# Moving beyond the motor cortex: a brain-wide evaluation of target locations for intracranial speech neuroprostheses

**DOI:** 10.1101/2024.11.29.626019

**Authors:** Maxime Verwoert, Maarten C. Ottenhoff, Simon Tousseyn, Johannes P. van Dijk, Pieter L. Kubben, Christian Herff

**Affiliations:** Department of Neurosurgery, Mental Health and Neuroscience Research Institute, Maastricht, The Netherlands; Academic Center for Epileptology, Kempenhaeghe/Maastricht University Medical Center, Heeze, The Netherlands; Department of Orthodontics, Ulm University, Ulm, Germany; Department of Electrical Engineering, Eindhoven University of Technology, Eindhoven, The Netherlands; Academic Center for Epileptology, Kempenhaeghe/Maastricht University Medical Center, Maastricht, The Netherlands

## Abstract

Speech is the fastest and most natural form of communication, which can be impaired in certain disorders. Speech brain-computer interfaces (BCIs) offer a solution by decoding brain activity into speech. Current neuroprosthetic devices focus on the motor cortex, which might not be usable in all patient populations. Fortunately, many other brain regions have been associated with the speech production process. Here, we investigate which regions are potential (alternative) targets for a speech BCI across a brain-wide distribution within a single study. The distribution includes sulci and subcortical areas, sampled with both a high temporal and a high spatial resolution. Thirty participants were recorded with intracranial electroencephalography during speech production, resulting in 3249 recorded contacts across the brain. We trained machine learning models to continuously predict speech from a brain-wide global to a single-channel local scale. Within each scale we examined a variation of selected electrode contacts based on anatomical features within participants. We found significant speech detection in both gray and white matter tissue, no significant difference between gyri and sulci at any of the analysis scales and limited contribution from subcortical areas. The best potential targets in terms of decoding accuracy and consistency are located within the depth of and surrounding the lateral fissure bilaterally, such as the (sub)central sulcus, transverse temporal gyrus (Heschls’ gyrus), the supramarginal cortex and parts of the insula. These results highlight the potential benefits of extending beyond the motor cortex and reaching the sulcal depth for speech neuroprostheses.

## Introduction

Decoding neural signals to restore natural and efficient communication for individuals with speech impairments is a key objective of speech brain-computer interface (BCI) research^1^. Speech production is a highly complex process involving an extensive and interconnected network of brain regions^2,3^. Yet, the current state-of-the-art speech BCIs are primarily focused on decoding signals from the sensorimotor cortex^4–9^. While this region has shown promise in enabling communication for patients with dysarthria or anarthria due to amyotrophic lateral sclerosis (ALS) or a brain-stem stroke, it may not cover the full intent of the user^10^ nor translate to individuals with different diseases or disease progressions. This is especially the case for individuals with damage to the motor cortex, which may happen due to a stroke or degeneration of cortical motor neurons in ALS^11^.

Given the extensive network of brain areas contributing to produce speech, it is essential to consider alternative regions across the entire brain. Existing non-invasive techniques that can capture signals across the brain, such as functional magnetic resonance imaging (fMRI), scalp electro-encephalography (EEG), and magneto-enecephalography (MEG), lack the simultaneous temporal and spatial resolution necessary to capture the rapid and complex dynamics of natural speech production for a continuous BCI. The intracranial technologies electro-corticography (ECoG) and micro-electrode arrays (MEAs) provide high temporal-spatial fidelity and have demonstrated success in speech BCIs^4–6,9^. However, both cover only the brain’s outer surface in typically a few specific cortical regions (i.e., the sensorimotor cortex). While research with these technologies provide crucial insights, these tools cannot access deeper areas of the brain—such as the sulci, inter-hemispheric regions, and subcortical nuclei.

Stereo-electroencephalography (sEEG), in contrast, offers an opportunity to probe these deeper areas of the brain. Research utilizing sEEG has revealed involvement of subcortical structures in speech production^12,13^, as well as cortical areas nestled within the sulci that are beyond the reach of surface-based recording technologies^14,15^. An added advantage of sEEG is the typical broad placement of electrodes across multiple regions, often bilaterally, within a single individual^16^, enabling a more diverse survey of brain regions implicated in speech. Although sEEG sampling can be spatially sparse, by aggregating data across multiple participants with varying electrode placements, we can achieve a brain-wide investigation, as we demonstrate in the current study.

Different representations of speech (e.g., articulatory, acoustic or semantic components) can be decoded from varying locations^17,18^, which may be beneficial for individuals with diverse brain injuries affecting speech. For instance, in motor-based impairments like apraxia of speech, where motor planning is disrupted^19^, BCIs may benefit from targeting areas associated with higher-level phonological or acoustic representations, such as the posterior superior temporal cortex^20–22^. In this study, we apply a continuous speech/silence decoder to 3249 widely distributed recording sites, covering gyri, sulci and deeper structures to investigate potential targets for speech neuroprostheses.

## Methods

### Participants

We recorded data from 30 individuals with medication-resistant epilepsy in the current study (Table 1). The data from the first 10 participants (P01 to P10) have previously been published^23^. All participants joined the study voluntarily and signed written informed consent. The study has been approved by the Institutional Review Boards of both Maastricht Unviversity (METC 2018-0451) and Epilepsy Center Kempenhaeghe. Participants were native Dutch speakers and were implanted with sEEG electrodes (Fig. 1A) to localize the epileptic onset zone. Electrode placement was solely determined based on clinical needs.

**Table 1.**
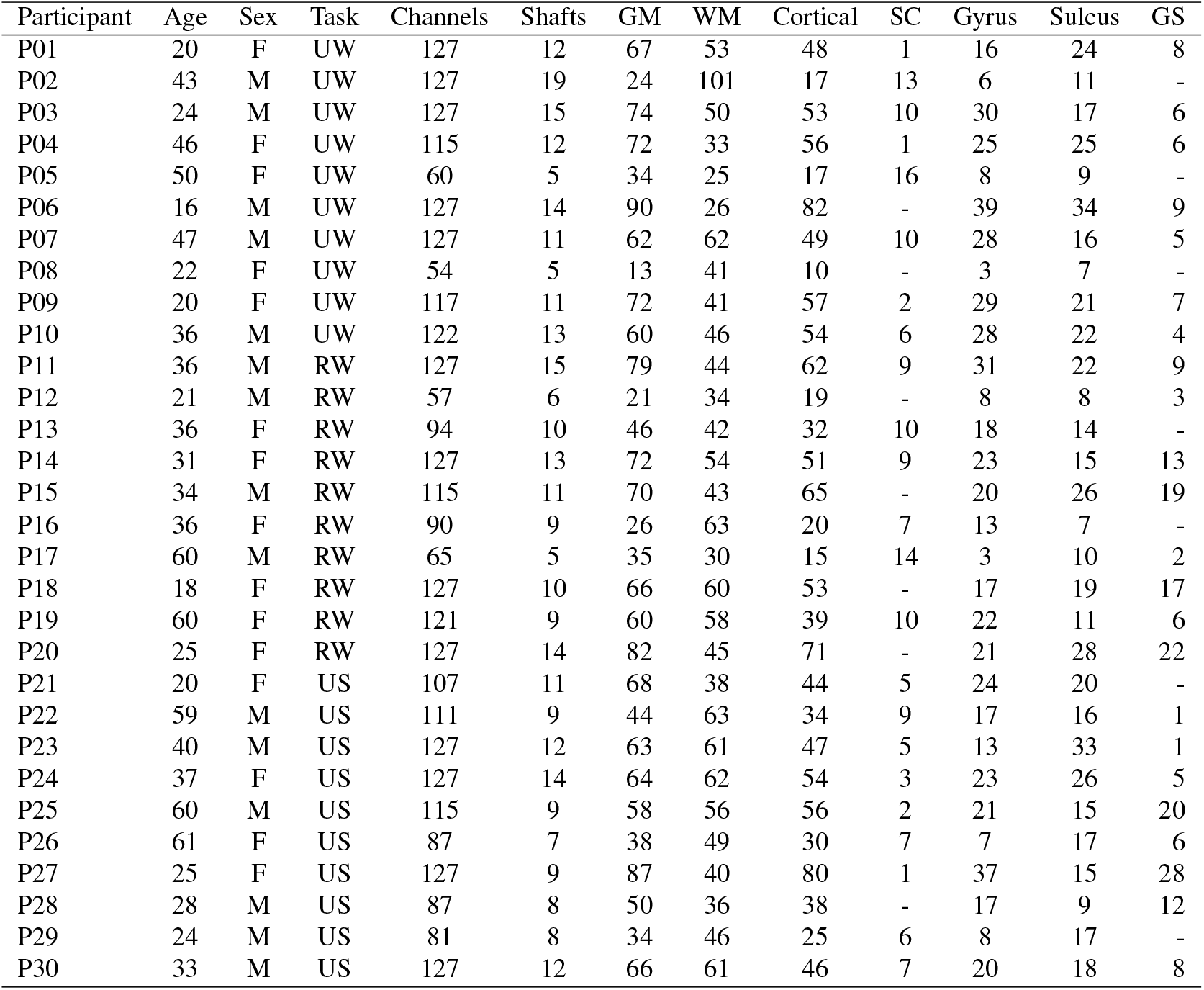
Demographic information, the performed task(s), the total number of channels and shafts that were recorded and the total number of channels available in the electrode selection categories for each participant. M = male; F = female; UW = unique words, RW = repeated words, US = unique sentences, GM = gray matter, WM = white matter, SC = subcortical, GS = gyrus-sulcus.

| Participant | Age | Sex | Task | Channels | Shafts | GM | WM | Cortical | SC | Gyrus | Sulcus | GS |
| --- | --- | --- | --- | --- | --- | --- | --- | --- | --- | --- | --- | --- |
| P01 | 20 | F | UW | 127 | 12 | 67 | 53 | 48 | 1 | 16 | 24 | 8 |
| P02 | 43 | M | UW | 127 | 19 | 24 | 101 | 17 | 13 | 6 | 11 | - |
| P03 | 24 | M | UW | 127 | 15 | 74 | 50 | 53 | 10 | 30 | 17 | 6 |
| P04 | 46 | F | UW | 115 | 12 | 72 | 33 | 56 | 1 | 25 | 25 | 6 |
| P05 | 50 | F | UW | 60 | 5 | 34 | 25 | 17 | 16 | 8 | 9 | - |
| P06 | 16 | M | UW | 127 | 14 | 90 | 26 | 82 | - | 39 | 34 | 9 |
| P07 | 47 | M | UW | 127 | 11 | 62 | 62 | 49 | 10 | 28 | 16 | 5 |
| P08 | 22 | F | UW | 54 | 5 | 13 | 41 | 10 | - | 3 | 7 | - |
| P09 | 20 | F | UW | 117 | 11 | 72 | 41 | 57 | 2 | 29 | 21 | 7 |
| P10 | 36 | M | UW | 122 | 13 | 60 | 46 | 54 | 6 | 28 | 22 | 4 |
| P11 | 36 | M | RW | 127 | 15 | 79 | 44 | 62 | 9 | 31 | 22 | 9 |
| P12 | 21 | M | RW | 57 | 6 | 21 | 34 | 19 | - | 8 | 8 | 3 |
| P13 | 36 | F | RW | 94 | 10 | 46 | 42 | 32 | 10 | 18 | 14 | - |
| P14 | 31 | F | RW | 127 | 13 | 72 | 54 | 51 | 9 | 23 | 15 | 13 |
| P15 | 34 | M | RW | 115 | 11 | 70 | 43 | 65 | - | 20 | 26 | 19 |
| P16 | 36 | F | RW | 90 | 9 | 26 | 63 | 20 | 7 | 13 | 7 | - |
| P17 | 60 | M | RW | 65 | 5 | 35 | 30 | 15 | 14 | 3 | 10 | 2 |
| P18 | 18 | F | RW | 127 | 10 | 66 | 60 | 53 | - | 17 | 19 | 17 |
| P19 | 60 | F | RW | 121 | 9 | 60 | 58 | 39 | 10 | 22 | 11 | 6 |
| P20 | 25 | F | RW | 127 | 14 | 82 | 45 | 71 | - | 21 | 28 | 22 |
| P21 | 20 | F | US | 107 | 11 | 68 | 38 | 44 | 5 | 24 | 20 | - |
| P22 | 59 | M | US | 111 | 9 | 44 | 63 | 34 | 9 | 17 | 16 | 1 |
| P23 | 40 | M | US | 127 | 12 | 63 | 61 | 47 | 5 | 13 | 33 | 1 |
| P24 | 37 | F | US | 127 | 14 | 64 | 62 | 54 | 3 | 23 | 26 | 5 |
| P25 | 60 | M | US | 115 | 9 | 58 | 56 | 56 | 2 | 21 | 15 | 20 |
| P26 | 61 | F | US | 87 | 7 | 38 | 49 | 30 | 7 | 7 | 17 | 6 |
| P27 | 25 | F | US | 127 | 9 | 87 | 40 | 80 | 1 | 37 | 15 | 28 |
| P28 | 28 | M | US | 87 | 8 | 50 | 36 | 38 | - | 17 | 9 | 12 |
| P29 | 24 | M | US | 81 | 8 | 34 | 46 | 25 | 6 | 8 | 17 | - |
| P30 | 33 | M | US | 127 | 12 | 66 | 61 | 46 | 7 | 20 | 18 | 8 |

**Figure 1.**
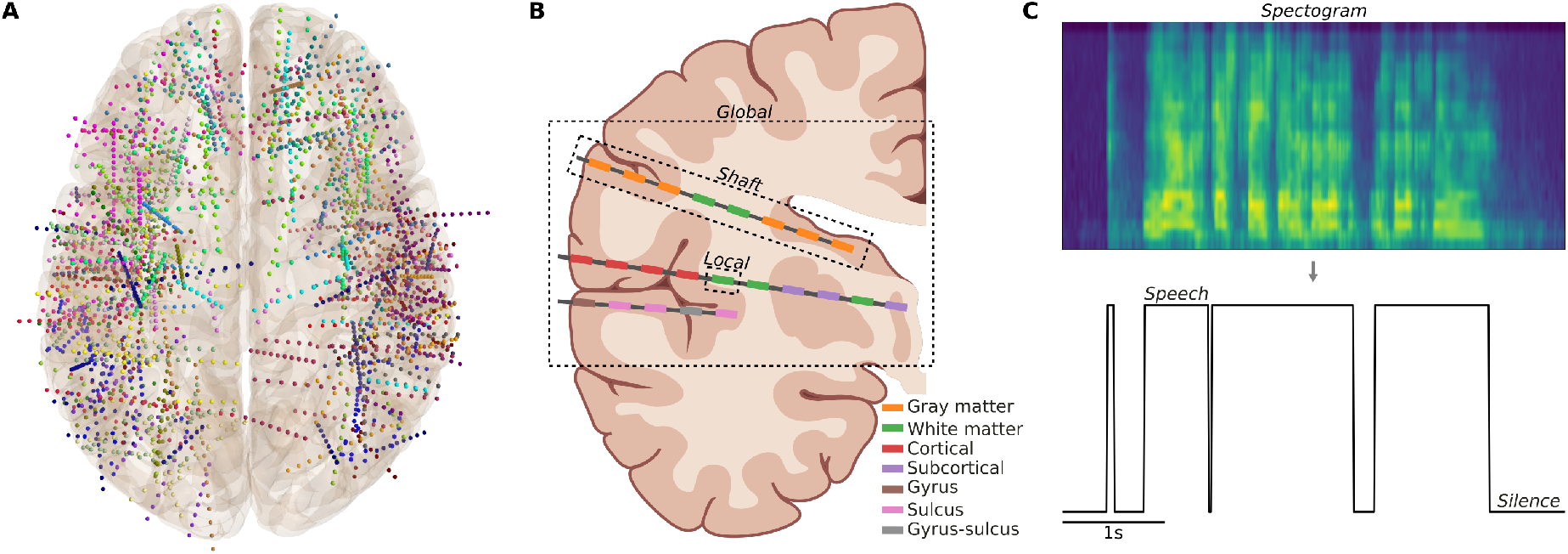
Graphical overview of the methods. **A)** Locations of recorded electrode contacts in an averaged brain from a top view. Colors differentiate the 30 participants. **B)** Graphical illustration of the analysis scales and electrode selection categories within a coronal brain slice. The ‘all’ category is not displayed as it includes all contacts and no specific selection. **C)** Example of how the spectrogram of the recording speech audio was used to extract the ‘speech’ and ‘silence’ labels used for the decoders.

### Tasks

Participants performed one of three speech production tasks in which they spoke 100 Dutch words or sentences out loud. In the SWPD dataset^24^ (P01 to P10), unique words (UW) were presented for two seconds. The next set of participants (P11 to P20) spoke 20 words, in random order, that were chosen to be relevant for patients in locked-in state^25^. The block of 20 words was repeated 5 times (RW). The last set of participants (P21 to P30) were presented with unique sentences (US). Since the reading speed varies between people, the first five participants of this set (P21 to P26) chose beforehand how much time they would like for reading each sentence. For the last five participants we modified the task design to allow the experimenter to press a key to end the speech trial once the participant finished reading the sentence. Across all tasks, each speech trial was followed by a 1-second rest period.

### Data acquisition

The neural data was recorded with one or two Micromed SD LTM amplifiers (Micromed S.p.A., Treviso, Italy) of 64 channels each, with a sampling rate of either 1024 Hz or 2048 Hz. The 2048 Hz signal was subsequently downsampled to 1024 Hz. The sEEG electrode shafts (Microdeep intracerebral electrodes; Dixi Medical, Beçanson, France) had a diameter of 0.8 mm, contact length of 2 mm and an inter-contact distance of 1.5 mm. The number of contacts on an electrode shaft could vary between 5 and 18. The number of implanted shafts varied between 5 and 19. The audio data was recorded with the microphone of the recording laptop (HP Probook) with a sampling rate of 48 kHz. LabStreamingLayer^26^ and our in-house software T-Rex^27^ was used to synchronize neural, audio and stimulus data.

### Electrode localization

The electrodes were localized for each participant with the img_pipe^28^ Python package and Freesurfer. The pipeline included the co-registration between a pre-implantation T1-weighted anatomical magnetic resonance imaging (MRI) scan and a post-implantation computerized tomography (CT) scan, manual identification of two contacts from each electrode, a linear inter- and/or extrapolation of the contacts accounting for the inter-electrode distance, a manual inspection of each contacts location and a non-linear warping to an average brain (MNI152) for visualisations only. The native MRI scan was segmented using the Fischl atlas^29^ to extract subcortical labels and parcellated using the Destrieux atlas^30^ to extract cortical labels for each contact. There were 3807 implanted electrode contacts of which 3249 were recorded. From here on, we only focus on the recorded contacts. There were 140 unique labels, 1419 had the ‘Cerebral White Matter’ label (43.67%) and 318 the ‘Unknown’ label (9.79%). The ‘Unknown’ label was given to contacts outside of brain tissue. The remaining 46.54% of contacts were spread across a variety of locations, including cortical and subcortical regions.

### Re-referencing

The data is recorded with a white matter reference (WMR), a contact located in white matter that is hand-picked by the treating neurologist as a suitable reference contact. It is usually a channel with a low amplitude signal that does not show epileptic activity. However, the use of one single contact for a reference can bias the signal in other contacts, solely due to spatial differences. Moreover, the reference contact would need to be located on the implantation shaft itself for a simulation of the performance of a single shaft. Thus, we applied an electrode shaft re-reference (ESR) by subtracting the average signal from all contacts within the same shaft from each channel. This is similar to the commonly used common average reference, except that it is restricted to the same electrode shaft rather than all implanted contacts across the brain. It is noteworthy that, in most cases, there were more electrodes implanted than recorded due to a limited number of channels in the amplifiers. Thus, the neurologist had to select which contacts to record data from. Sometimes this lead to missing contacts within a shaft of electrodes.

### Signal processing

The neural signals were filtered to the broadband high-frequency activity (70-170 Hz) with an IIR bandpass filter (filter order 4) and the Hilbert envelope was extracted. Two IIR bandstop filters (filter order 4) were applied to attenuate the first two harmonics of the 50 Hz line noise. The filters were employed forward and backward to eliminate a potential phase-shift. The envelope was averaged in 50 ms windows with a 10 ms frameshift as the neural features. The audio signal was downsampled to 16 kHz and the Short-Term-Fourier-Transform was applied to extract audio features, also in 50 windows with a 10 ms frameshift. This procedure ensured an alignment between the labels and the neural features. An energy threshold was calculated with the maximum plus minimum value of the average energy across spectral bins, multiplied by a static value of 0.45. This threshold was applied to the audio features to extract ‘speech’ and ‘silence’ labels.

### Analysis scales

The data was analyzed at three different scales. In the ‘global’ scale, all recorded electrode contacts within an individual were included as features in the model. In the ‘shaft’ scale, the analysis was restricted to all contacts within a shaft. Finally, in the ‘local’ scale, each contact was analyzed individually (Fig. 1B).

### Electrode selection categories

For each of the analysis scales, we additionally selectively included contacts located in specific tissue types or structures in separate analyses to determine the contribution of each. Due to a margin of error in electrode labeling, we utilized the proximal tissue density (PTD) for each contact to determine if it was located in ‘white matter’ or ‘gray matter’. The PTD score indicates the relative proportion of gray versus white matter in the 26 MRI voxels surrounding the center of the contact^31^, using the following formula:

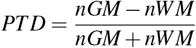

where *nGM* and *nWM* represent the number of gray matter and white matter voxels, respectively. We included subcortical areas and labeled these as gray matter. For the white matter analysis, we only included contacts with *PTD* < 0. For the gray matter analysis, we only included contacts with *PTD* > 0. The PTD could not be calculated for some contacts as they were surrounded entirely by non-brain tissue, these were excluded in both analyses.

Next, we used the labels from the atlases to further sort the contacts into the categories ‘cortical’, ‘subcortical’, ‘gyrus’, ‘sulcus’ and ‘gyrus-sulcus’. The ‘cortical’ category refers to those contacts labeled by the atlas as gray matter. However, these do not include the subcortical structures as they do in the general ‘gray matter’ analysis. In this case, the subcortical structures (e.g., the hippocampus) are labeled as ‘subcortical’. The ‘gyrus’ category refers to contacts located near the cortical surface (e.g., the precentral gyrus), whereas the ‘sulcus’ refers to contacts located deeper within a cortical fold (e.g., the central sulcus) which could not be reached with a standard electrode grid. The ‘gyrus-sulcus’ category refers to contacts located on the border between a gyrus and sulcus, which could not be clearly separated and therefore have this combined label in the atlas (e.g., the subcentral gyrus and sulcus). Note that some regions within the lateral fissure are gyral, even when they cannot be reached with electrode grids (e.g., the short insular gyrus).

### Speech detection

Neural features were enhanced with causal temporal information from non-overlapping windows up to 200 ms into the past. This means that for each channel, there were 4 additional neural features with a shifted time alignment to the labels. Only temporal windows from the past were used, as future windows could not be used in a real-time BCI. The data was split into a 10-fold cross-validation. Neural features were normalized to zero mean and unit variance using the mean and standard deviation of the training data. The same normalization was then applied to the testing data.

We employed a continuous binary classification approach to distinguish between speech and silence. This simple framework facilitates analysis across tasks and serves as a foundation for a basic BCI. The goal was not to optimize classification performance but rather to identify potential brain areas involved in speech production. For this, we utilized a Linear Discriminant Analysis (LDA) classifier. Given the class imbalances, with generally more silence than speech, we used balanced accuracy as the primary outcome metric to account for this disparity.

This analysis was computed for each of the electrode selection categories described in the previous section. Additionally, the single channel analysis was repeated for each individual temporal window from −500 ms up until +500 ms to investigate temporal dynamics.

### Significance threshold

We computed a distribution of chance level accuracy scores by randomly shuffling the speech/silence labels, calculating the balanced accuracy between the original and shuffled labels in 10 folds similar to the speech detection pipeline, and repeating this procedure 1,000 times for each participant. Next, we calculated the 99th percentile (*α* = 0.01) within each fold per participant and set the individual threshold at the maximum across folds. As the final significance threshold, we took the maximum score across participants. This procedure led to a strict threshold of 52.44%, well above the theoretical chance level of 50%.

### Regions-of-interest

Due to variations in sampling within and between regions as they are labeled by the atlases, we grouped results together in 9 larger regions-of-interest (ROIs) across the brain. A detailed list of the anatomical regions from the atlases included in each ROI is provided in the supplementary material (Table S1). Note that only the subcortical regions that were sampled were included in the list.

## Results

In this work, we recorded speech production sEEG data from 30 participants with electrode contacts covering a wide range of locations (Fig. 1A) and structures/tissue types (Fig. 1B). We used a binary classification approach to decode the recorded neural activity into speech or silence labels (Fig. 1C). Electrodes were analyzed at three levels (global, shaft and local), with additional analyses discriminating tissue types and brain structures (Fig. 1B).

## Global to local scale

Speech can be detected well above chance level for all 30 individuals (ranging between 60.13% and 93.79%) at the global scale using all recorded contacts (Fig. 2A). This holds true when only including contacts in gray matter as well. The accuracy of the results dropped slightly for certain individuals in the other categories and the performance goes down to below significance threshold for most individuals using only subcortical contacts. It is important to note that the lowest performing categories, subcortical and gyrus-sulcus, also had a much lower number of channels within individuals compared to the other categories (Table 1). Independent samples t-test were performed to compare certain electrode selection categories with one another. Gray matter only was not significantly different from white matter only (*t*(58) = 1.47, *p* = 0.15), nor was the sulcus significantly lower than the gyrus category (*t*(58) = 0.67, *p* = 0.51). However, the cortical category was higher than the subcortical one (*t*(51) = 9.20, *p* < 0.001) and both the gyrus (*t*(51) = 3.25, *p* = 0.002) and the sulcus (*t*(51) = 2.49, *p* = 0.016) scored higher than the gyrus-sulcus category. The variation in performance between individuals could be related to the sheer amount of features included, as there was a significant correlation (Pearson’s *r*(58) = 0.43, *p* = 0.018) between the amount of features and accuracy (Fig. 2B), and/or the specific anatomical regions that were sampled.

**Figure 2.**
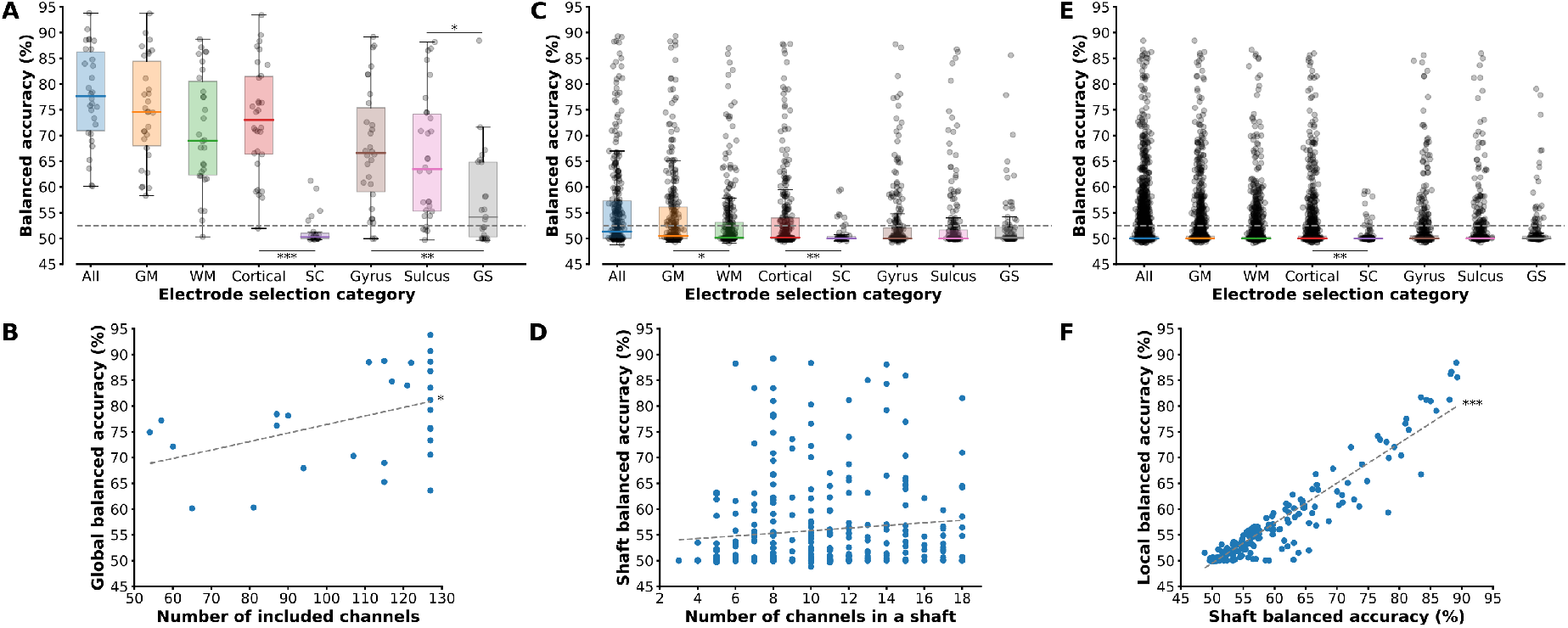
Distribution of balanced accuracy results within electrode selection categories across the analysis scales and correlations. **A)** Global scale balanced accuracy results. **B)** Correlation between global accuracy and number of features. **C)** Shaft scale balanced accuracy results. **D)** Correlation between shaft accuracy and shaft size. **E)** Local scale balanced accuracy results. **F)** Correlation between maximum channel accuracy within a shaft and accuracy of the complete shaft. In A-C-E the individual dots represent the averaged balanced accuracy per individual, shaft or channel, respectively. The box plots show the median and inter-quartile range, whiskers show the maximum range excluding outliers. The dashed gray line represents the significance threshold in A-C-E and the best-fit linear trend, calculated using a least-squares linear regression, in B-D-F. ^*^*p* < 0.05, ^**^*p* < 0.01, ^***^*p* < 0.001.

At the shaft scale, there are large variations between individual shaft results with the median accuracy being below the significance threshold in all of the categories (Fig. 2C). However, the maximum scores (89, 89, 87, 88, 59, 88, 87 and 86%) are not much lower than in the global scale (94, 94, 89, 93, 61, 89, 88 and 88%). At this level, between shafts overall, there was a significant difference between the gray and white matter categories (*t*(602) = 2.25, *p* = 0.025) and between the cortical and subcortical ones (*t*(352) = 3.00, *p* = 0.003). There was no significant difference between the gyrus and sulcus (*t*(464) = 0.23, *p* = 0.82), nor either with the gyrus-sulcus category (gyrus: *t*(329) = 0.29, *p* = 0.77; sulcus: *t*(295) = 0.12, *p* = 0.91). At the shaft level, there was not a significant correlation (Pearson’s *r*(634) = 0.10, *p* = 0.07) between the size of the shaft and accuracy (Fig. 2D), suggesting that the anatomical location of the shaft may be a more important factor than the sheer amount of features included.

In the local scale, the results are similar those at the shaft scale, with large variations between channels within categories and most channels below the significance threshold (Fig. 2E). However, the maximum scores of single channels (88, 88, 87. 86, 59, 86, 86 and 79%) are still similar to those in the shaft and global scales. In these large groups, there are no statistically significant differences between gray matter and white matter (*t*(3170) = 1.95, *p* = 0.05), between the gyrus and sulcus (*t*(1105) = 0.28, *p* = 0.78), the gyrus versus gyrus-sulcus (*t*(790) = 1.22, *p* = 0.22), nor the sulcus versus gyrus-sulcus (*t*(747) = 1.00, *p* = 0.32). The subcortical category is significantly lower than the cortical category (*t*(1485) = 3.06, *p* = 0.002). There was a strong correlation (Pearson’s *r*(634) = 0.95, *p* < 0.001) between the maximum accuracy of a single channel within a shaft and the accuracy of the complete shaft (Fig. 2F), suggesting that one or a few channels within a shaft are driving most of the results.

### Anatomical contributions

We zoom into the local scale as particular anatomical regions may be more important than the generic categories presented above. Since the global and shaft scales cover multiple regions at once, we can only compare regions at the local channel level. In Figure 3A we see the classification accuracy distribution across all significant channels. This figure indicates that the best decoding results are clustered within and around the lateral fissure, with a more focal clustering in the right hemisphere.

**Figure 3.**
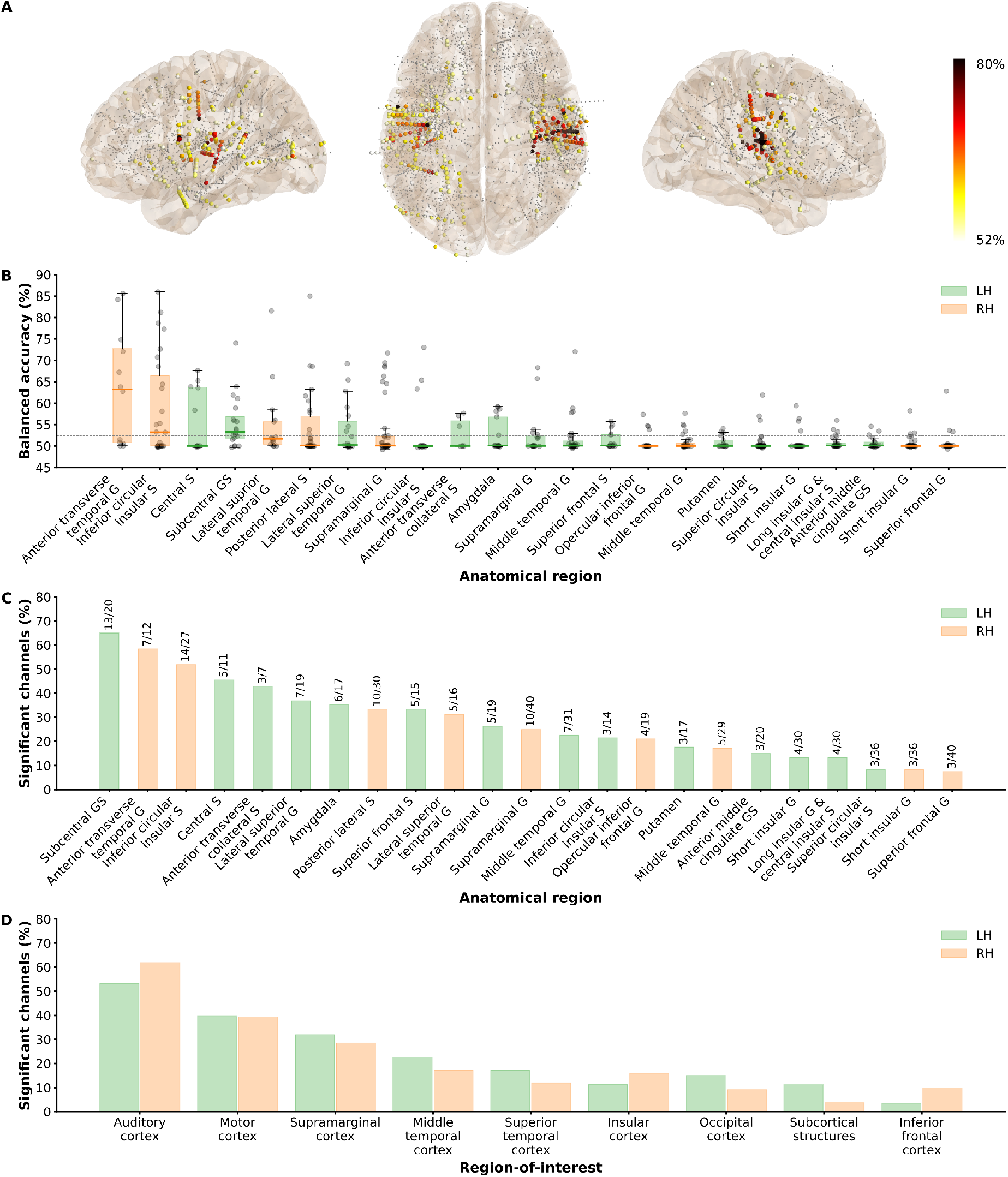
Anatomical dissemination at the local scale. **A)** Spatial distribution of significant balanced accuracy scores across channels in an averaged brain. The colorbar ranges from the significance threshold (52%) to ≥ 80%. Smaller gray dots represent insignificant channels. **B)** Distribution of balanced accuracy scores within anatomical regions, sorted by average accuracies. The color of the bar indicates the hemisphere. **C)** Percentage of significant channels within anatomical regions, sorted by percentage. The text above the bar describes the amount of significant and total amount of channels within that region. The color of the bar indicates the hemisphere. **D)** Percentage of significant channels within larger regions-of-interest within each hemisphere, sorted by the average percentage between the two hemispheres. G = gyrus, S = sulcus, GS = gyrus-sulcus.

Out of the 137 sampled anatomical regions (excluding white matter and unknown labels), we further focus on the 23 regions that were sampled by at least 5 different participants and had at least 3 significant channels. The right anterior transverse temporal gyrus (Heschl’s gyrus) scored the highest in terms of accuracy (Fig. 3B) and second in terms of consistency (the percentage of significant channels within that region, Fig. 3C), the left Heschl’s gyrus was not sampled enough. The right inferior circular insula scored the second highest in accuracy and third in consistency, while only at the 9th and 14th place in the left hemisphere. The left central sulcus and subcentral gyrus-sulcus (GS) both scored high in accuracy and consistency, with over 60% significant channels in the subcentral GS. These regions, along with the precentral gyrus, were not sampled by enough participants in the right hemisphere. However, the left precentral gyrus was, and only had 2/12 significant contacts. When we look at the larger regions-of-interests (ROIs; Fig. 3D), including regions not sampled enough individually, we see that generally the motor cortex was equally involved between the hemispheres.

We further see bilateral regions surrounding the ones described above, such as the superior and middle temporal cortex, the supramarginal cortex, the inferior frontal gyrus and other parts of the insula. The most consistent results overall come from the auditory, motor and supramarginal regions (Fig. 3D). Note that parts of the superior temporal cortex are incorporated in the auditory cortex ROI, the remaining parts of the relative large superior temporal cortex ROI are therefore not very consistently involved. The insular ROI follows, as it was particularly the inferior part that scored high in accuracy and consistency. It is worth noting that, while we look at the relative number of significant channels, the channels were not sampled exactly uniformly within regions and between hemispheres, which limits the interpretability of comparisons.

### Temporal dynamics

We explored how the spatial distribution and number of significant channels varied between temporally misaligned features and tasks (Fig. 4). The results were grouped in five larger time segments (Fig. 4A-E), representing time windows well before alignment (−500 until −300 ms), just prior to alignment (−300 until −100 ms), surrounding alignment (−100 until +100 ms), just after alignment (+100 until +300 ms) and well after alignment (+300 until +500 ms). Neural features just prior to alignment resulted in the largest number of channels significantly detecting speech (Fig. 4F), with a steeper drop-off after than before alignment. The variation between tasks reflects the variation in the length of the speech trials (short words versus long sentences). All three tasks engaged peri-sylvian, temporal and sensorimotor areas, while only the sentences (US) task engaged frontal areas.

**Figure 4.**
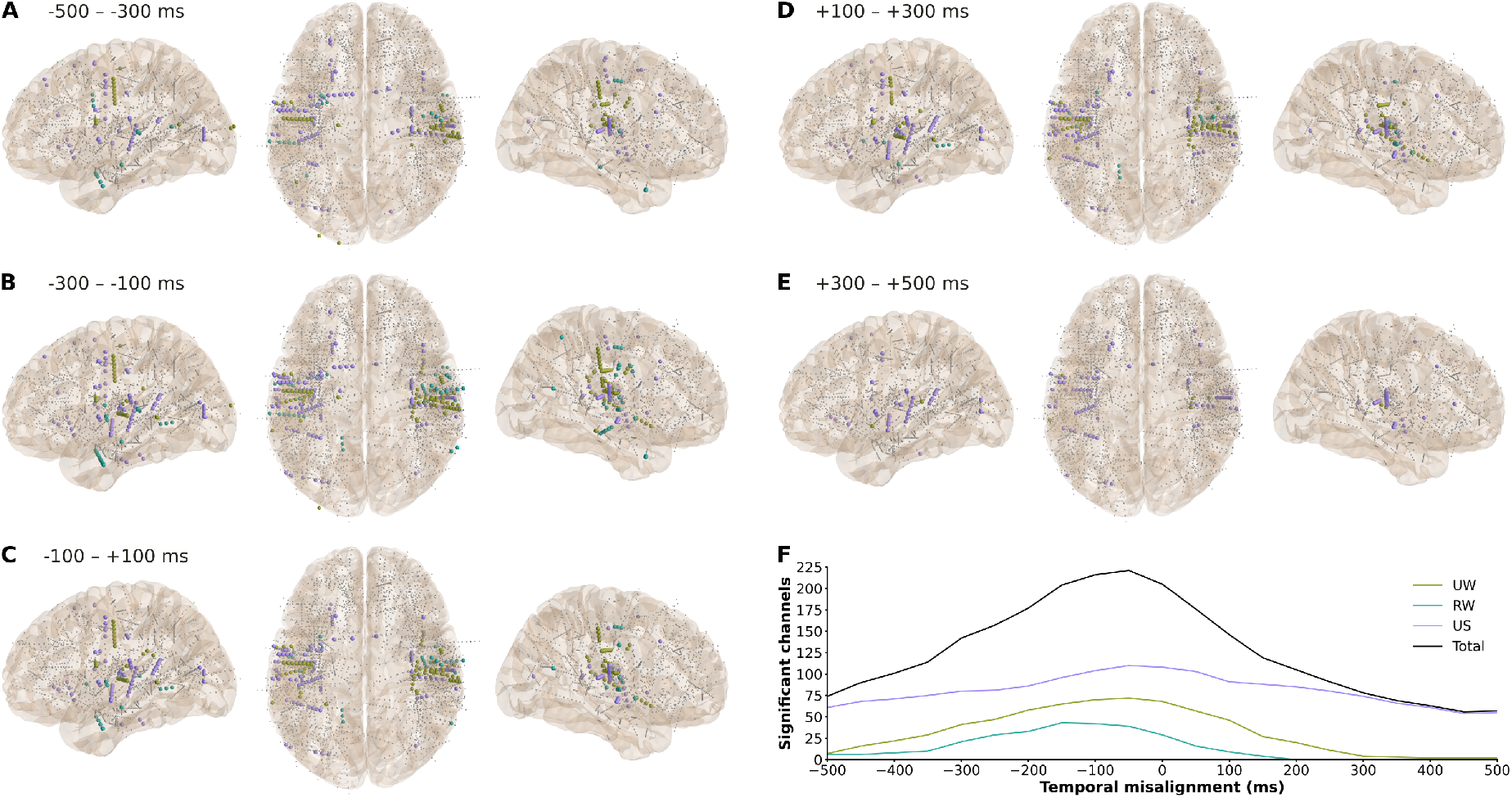
Temporal dynamics. **A-E)** Significant channel locations with colors representing the three tasks, divided into five time segments. Smaller gray dots represent non-significant channels. **F)** Number of significant channels across time-frames from all (brain-wide) contacts. UW = unique words task, RW = repeated words task, US = unique sentences task.

## Discussion

Speech detection was performed at three scales of analysis and eight electrode selection categories. Speech was reliably detected for all thirty individuals using all available data within individuals (global scale), but the performance varied greatly between electrode shafts (shaft scale) and individual channels (local scale). While at a global scale the performance was correlated with the amount of included channels, this was not the case at the shaft scale, but instead correlated strongly with the accuracy of the best individual channel along the shaft. These results combined suggest that the location of individuals channels is important, while a combination of multiple distributed regions may be beneficial as well. Previous research has indeed showed that overtly and covertly produced speech can be reconstructed well from distributed regions in offline^32,33^ and in real-time^33^ decoding.

Overall, the gray matter category scored slightly higher than the white matter category at the shaft scale. At the global scale the difference was not significant likely due to difference in the number of data-points. Whilst lower than gray matter, the nonetheless high accuracy results from contacts located in white matter indicates that we may be able to decode valuable information directly from white matter tracts. This activity may reflect volume conduction from a combination of nearby and distant gray matter regions^31^. For example, direct electrical stimulation of the anterior arcuate fasciculus tract, superior to the insula, has been found to induce thoughts or alter the conscious awareness of thoughts, similar to stimulation of the posterior parietal cortex itself^34^. Our results are in line with previous decoding studies with sEEG that have also found contributions from white matter contacts^15,35,36^. However, in our work, these findings could, at least in part, have been due to spreading of information from cortical contacts within the same electrode shaft due to re-referencing. It is also a challenge to locate the exact white matter tracts, requiring additional anatomical analyses and/or diffusion-tensor imaging, and therefore beyond the scope of our current work.

There were clear differences between the cortical and subcortical categories in all analysis scales. While there were a few significant channels in subcortical areas, the accuracy did not reach nearly as high as in cortical and even white matter regions. Previous studies have found language-related functions in the hippocampus^12,13,37,38^ and basal ganglia^39^ and other subcortical areas through a large meta-analysis^40^. Studies reveal specialized functions of these areas, such as semantic integration^37^ and syntactic processing^39^, and therefore may be too subtle for general speech detection as in the current study. Alternatively, in electrophysiology, we may need to investigate these areas with lower frequencies^37^ or even single cell recordings^13,41^.

Within the cortical channels, there was no statistically significant difference between the gyrus and sulcus categories at any of the analysis scales across the brain, despite general differences in cytoarchitecture and function^42^. Even within the local scale, only looking at significant contacts, we see both gyral and sulcus regions detecting speech. However, the gyri are more often mentioned in the speech decoding literature as sulci cannot be sampled with the commonly used electrocorticography grids. Note that, in this work, gyri do not necessarily mean they can be sampled with electrode grids on the surface of the brain, as they can be located within the lateral fissure. A previous sEEG study did analyze differences in speech detection accuracy across the level of depth along shafts and found generally higher accuracies at the surface level with overt and mouthed speech, but not with imagined speech^15^. These results suggest that the deeper regions may be just as important for an imaginary-based speech neuroprosthesis.

Next, we dived further into significant anatomical regions at the local scale. The overall spatial distribution across the brain corresponds with the core language network^2,3^ and its bilaterality is consistent with previous sEEG speech production studies^15,17,18,22^. At the top positions in terms of both accuracy and consistency are Heschl’s gyrus (the auditory cortex), the inferior insular sulcus and the (sub)central sulcus. While named gyrus, Heschl’s gyrus is also nestled deeper within the lateral fissure. Interestingly, the precentral gyrus (primary motor cortex) was sampled plenty in the left hemisphere, but only showed 2/12 significant contacts. This is in stark contrast with the sulcal parts of the motor cortex, suggesting that reaching the sulcal depth may be beneficial for motor-based BCIs^43^.

We further looked at larger regions-of-interest to generate a better overview across regions, including the individual areas that were left out of Figure 3B and C due to not being sampled by plenty of different participants or not containing plenty of significant contacts. Whilst sampling of the motor cortex was not ideal, the auditory areas of the superior temporal lobe showed the most consistent results, despite known suppression of regions within the auditory cortex during self-generated speech^44^. The supramarginal cortex may also be a good alternative for a speech neuroprosthesis, as was previously shown with a micro-electrode array even for imagined speech^45^. There was limited contribution from the inferior frontal cortex, which may be due to the reading task not requiring spontaneous speech planning^46^. The reading could also explain the relatively stronger occipital than frontal involvement overall. However, when we split the results between the three tasks for temporal dynamics, we do note that only the most complex (sentences) task reached significance in the frontal areas, likely engaging more higher-order executive control^47^ than single word reading.

As most of the significant regions are also known to be activated during speech or sound perception^3,48^, the question is whether they will remain activated without or with a limited amount of overt speech. Previous research has indeed found similar, albeit more attenuated, activation patterns with purely imaged speech, compared to overt or mouthed speech^15^. This even applies to auditory regions^15,49,50^ as they may be involved in the internal representation of speech^49^. However, responses to perceived speech would be another problem for a practical speech BCI. This is even the case for a speech neuroprosthesis on the motor cortex, with the potential of leading to false positives through speech perception^51,52^ or even through unintentional inner speech^52^. Ensuring executive control is an important ethical aspect in the development of speech neuroprostheses^53^ and efforts must therefore be made to mitigate false positives.

In the sensorimotor cortex, differential spatial patterns between produced speech, perceived speech and rest can be used as an ‘intended’ speech detector^51^. A similar spatial decoder may also be possible in the superior temporal cortex, considering differential patterns found between produced and perceived speech here as well^22,44^, with a gradient of more anterior regions preferring self-generated and posterior regions preferring perceived speech^22^. The hippocampus may also be interesting as it is found to have a stronger coupling with auditory cortex during self-generated speech versus hearing one’s own pre-recorded speech and is thought to have a predictive role in speech production^38^. However, we did not see hippocampal involvement in our current work. The posterior insula, similar to the area we see in this work, is also found to be more activated during self-generated than perceived speech^54^. However, the insula may be more related to the coordination of speech with autonomic functions and may therefore not be present during purely imagined speech^55^. In our earlier work, we have also shown that speech is better reconstructed from this region using neural features during speech articulation rather than before, unlike the other regions^17^. In our current work, while the largest amount of significant channels overall were based on neural features just prior to the speech features, we could not distinguish pre- and post-articulation signals due to the continuous nature of our analysis and different temporal dynamics between our speech tasks. Nonetheless, a speech detector from within or between such regions that can additionally differentiate between self-generated and perceived speech may be used to mitigate the false positive effect from speech perception, as well as as on/off switch that a participant can control to mitigate the potential effect from unintended inner thoughts^52,53^.

## Conclusion

This study provides a comprehensive overview of speech decoding throughout the brain, and underscores the potential of regions beyond the cortical surface for speech neuroprostheses. We showed that white matter contacts provided strong decoding accuracy. Sulci regions, which cannot be measured with ECoG or MEA, yielded decoding results on-par with gyral locations. Additional to the usually recorded left hemisphere, we found that locations on both hemispheres provided very high accuracies, with the most consistent results derived from auditory regions, followed by the motor and supramarginal cortices. Frontal areas provided significant decoding results only in the sentence task, indicating the importance of different types of speech in the training data. Overall, these findings indicate that there is large potential for speech neuroprostheses even when the motor cortex cannot be targeted.

## Data availability

The raw data is publicly available at https://doi.org/10.17605/OSF.IO/AK3DP and code can be found on https://github.com/neuralinterfacinglab/SpeechTargets.

## Acknowledgements

This publication is part of the project INTENSE (with project number 17619 of the research programme NWO Crossover Programme) which is (partly) financed by the Dutch Research Council (NWO). C.H. acknowledges funding by the Kavli Foundation.

## Author contributions statement

C.H. and P.K. and M.V. designed the experiments, C.H, M.C.O. and M.V. collected the data, M.V. ran the analyses, M.V. wrote the manuscript. All authors reviewed the manuscript and declare no competing interests.

## Supplementary Material

**Table S1.**
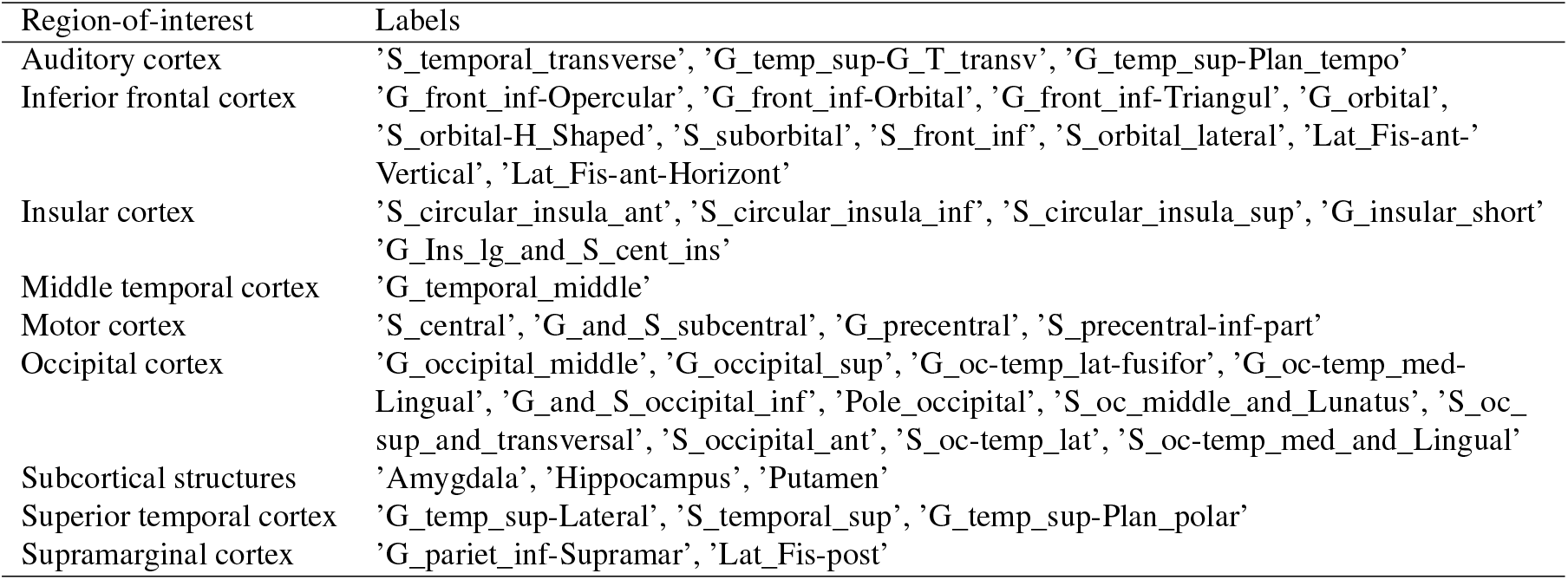
Included Destrieux/Fischl atlas labels per region-of-interest. The same labels were applied to both hemispheres.

